# Elucidating the genetic architecture of reproductive ageing in the Japanese population

**DOI:** 10.1101/182717

**Authors:** Momoko Horikoshi, Felix R. Day, Yoichiro Kamatani, Makoto Hirata, Masato Akiyama, Koichi Matsuda, Hollis Wright, Carlos A. Toro, Sergio R. Ojeda, Alejandro Lomniczi, Michiaki Kubo, Ken K. Ong, John R. B. Perry

## Abstract

Population studies over the past decade have successfully elucidated the genetic architecture of reproductive ageing. However, those studies were largely limited to European ancestries, restricting the generalizability of the findings and overlooking possible key genes poorly captured by common European genetic variation. Here, in up to 67,029 women of Japanese ancestry, we report 26 loci (all P<5×10¯^8^) for puberty timing or age at menopause, representing the first loci for reproductive ageing in any non-European population. Highlighted genes for menopause include *GNRH1,* which supports a primary, rather than passive, role for hypothalamic-pituitary GnRH signalling in the timing of menopause. For puberty timing, we demonstrate an aetiological role for receptor-like protein tyrosine phosphatases by combining evidence across population genetics and pre- and peri-pubertal changes in hypothalamic gene expression in rodent and primate models. Furthermore, our findings demonstrate widespread differences in allele frequencies and effect estimates between Japanese and European populations, highlighting the benefits and challenges of large-scale trans-ethnic approaches.

## Introduction

The first menstrual period (‘menarche’) and onset of menopause are key milestones of female reproductive ageing, representing the start and end of reproductive capacity. The timings of these events vary widely between individuals and are predictors of the rate of ageing in other body systems^1^. This variation reflects a complex mix of genetic and environmental factors that population-based studies are beginning to unravel. Over the past decade, successive waves of genome-wide association study (GWAS) meta-analyses have illuminated the genetic architecture of reproductive ageing and shed light on several underlying biological processes, many of which are also highlighted through studies of rare human disorders of reproduction^2-7^. Hence, puberty timing appears to be predominantly regulated by the central nervous system (CNS), including components of the hypothalamic-pituitary axis and complex molecular silencers of that system^5^. In contrast, the aetiological drivers of menopause timing appear to be ovary-centric, largely focussed on the ability of oocytes to maintain genome stability and hence preserve the ovarian primordial follicle pool^7^.

A key limitation of previous GWAS for reproductive ageing is their large circumscription to populations of European ancestry, due to the lack of available large-scale studies of other populations. This population restriction has limited the generalizability of the findings and may have led to a failure to detect key genes and pathways that are poorly represented by common functional variants in European populations. Previous small-scale genetic studies in East Asian and African-American samples have replicated a small number of the reported European loci^8,9^. However, such studies have not yet demonstrated any known or new genetic association signals for menarche or menopause timing at genome-wide statistical significance. To address this limitation, we performed two separate GWAS for ages at menarche and menopause in up to 67,029 women of Japanese ancestry from the BioBank Japan Project (BBJ)^10^. This sample represents a 3-fold larger sample size than any previous non-European ancestry GWAS. We identify 26 loci for ages at menarche or menopause at genome-wide significance, indicating several new genes and pathways. The analyses also revealed widespread differences in effect estimates between populations, highlighting the benefits and challenges of trans-ethnic GWAS meta-analyses.

## Results

### Effects of known European menopause and menarche loci in Japanese

Data on age at natural menopause and age at menarche were available on 43,861 and 67,029 genotyped women of Japanese ancestry, respectively. Genotyping array-based heritability in this Japanese sample was 10.4% (S.E. 0.9%) for menopause (contrasting with 36% in Europeans) and 13% (S.E. 0.6%) for menarche (32% in Europeans). Mean age at menarche in this Japanese population (overall: 13.9 years) was higher in than previously reported in contemporary European populations (12.4 to 13.7 years)^5^, but showed a marked secular trend in Japanese from 15.2 years in women born pre-1935 to 12.3 years in those born post-1965, and this was accompanied by increasing heritability (from 14.2% to 20.6%; P_het_=0.03, **Table S1**).

Of the 54 previously identified European menopause loci (**Table S2**), 52 were polymorphic in Japanese; of these 46 (88.4%) had a consistent direction of effect (binomial P=l×10¯^8^; 29 loci at nominal significance P<0.05). For menarche, 348/377 autosomal variants found in Europeans were present in the Japanese dataset (**Table S3**); of these 282/348 (81.0%) had a consistent direction of effect (binomial P=6.4×10¯^33^, 108 loci at P<0.05). In aggregate, genetic variation +/− 250kb from European-identified SNPs explained 2% (S.E. 0.2%) and 3.6% (S.E. 0.2%) of the trait variance for age at menopause and menarche, respectively (contrasting with 8.0% [S.E. 0.5%] and 8.4% [S.E. 0.4%] in Europeans, respectively).

There were notable differences in allele frequencies between populations at these European identified signals (**Table S2**), with 23 loci (2 menopause, 21 menarche) monomorphic in Japanese. The mean absolute difference in allele frequency was 17%, with the largest difference at the menarche locus, 20qll.21, where the C-allele at rs1737894 in Europeans (frequency ~60%) is absent in Japanese.

To compare effect sizes between populations at these previously identified signals, we calculated effect estimates for Europeans in up to 73,397 women from the UK Biobank study, independent of the European discovery samples. Across 52 (polymorphic in Japanese) menopause loci, 44 (84.6%) showed a larger effect in Europeans than in Japanese (binomial P=4×10¯^7^), 23 of which were significantly different (P_diff_ <0.05, **Table S2**). Similarly, for the 102 menarche loci that were previously identified in Europeans excluding UK Biobank (**Table S4**), 77 (75.5%) showed larger effects in UK Biobank Europeans than in Japanese (binomial P=2.5×10¯^7^, 22 at P_diff_ <0.05). These findings likely indicate widespread population differences in LD between GWAS signals and the underlying causal variants, or possible differences in modifying environmental factors.

### Novel menopause and menarche signals in Japanese

To identify novel genetic signals for ages at menopause and menarche in our Japanese population, we tested genome-wide markers imputed to the 1000 genomes Phase 3 reference panel^11^. For menopause, 16 independent signals reached genome-wide significance (P<5×10¯^8^), 8 of which are novel and not previously reported in Europeans (**Table 1; Figure 1**). Additionally, we found a novel signal (rs76498344, P_japanese_=3.6×10¯^12^) near the previously reported locus *MCM8* which was uncorrelated with the reported European lead SNP (rs451417, r^2^=0.03 with rs76498344 in Japanese) and showed no association in Europeans (rs76498344 P_Euro_=0.78)^5^. For menarche, 10 independent signals reached genome-wide significance, 2 of which are novel loci, and a third represents a novel Japanese-specific signal in a known European locus (r^2^~0) near *PTPRD* (**Table 1; Figure 1**).

**Figure 1.**
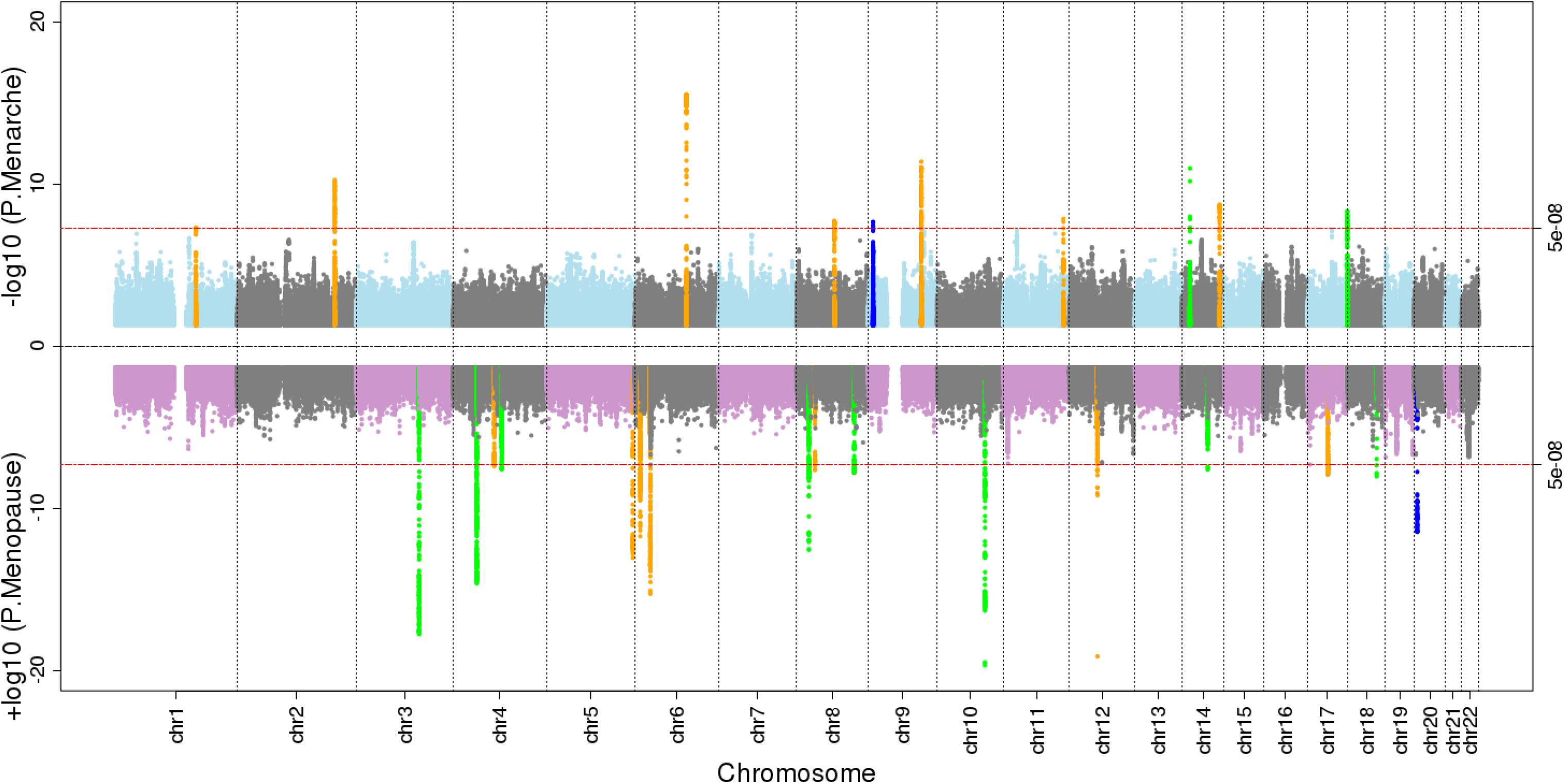
“Miami” plot showing genome-wide association test statistics for ages at menarche (top) and menopause (bottom) in the BioBank Japan Project. Genome-wide significant loci in BBJ are highlighted according to: novel loci (green), novel signals at known loci (blue) and known loci (orange)

**Table 1:**
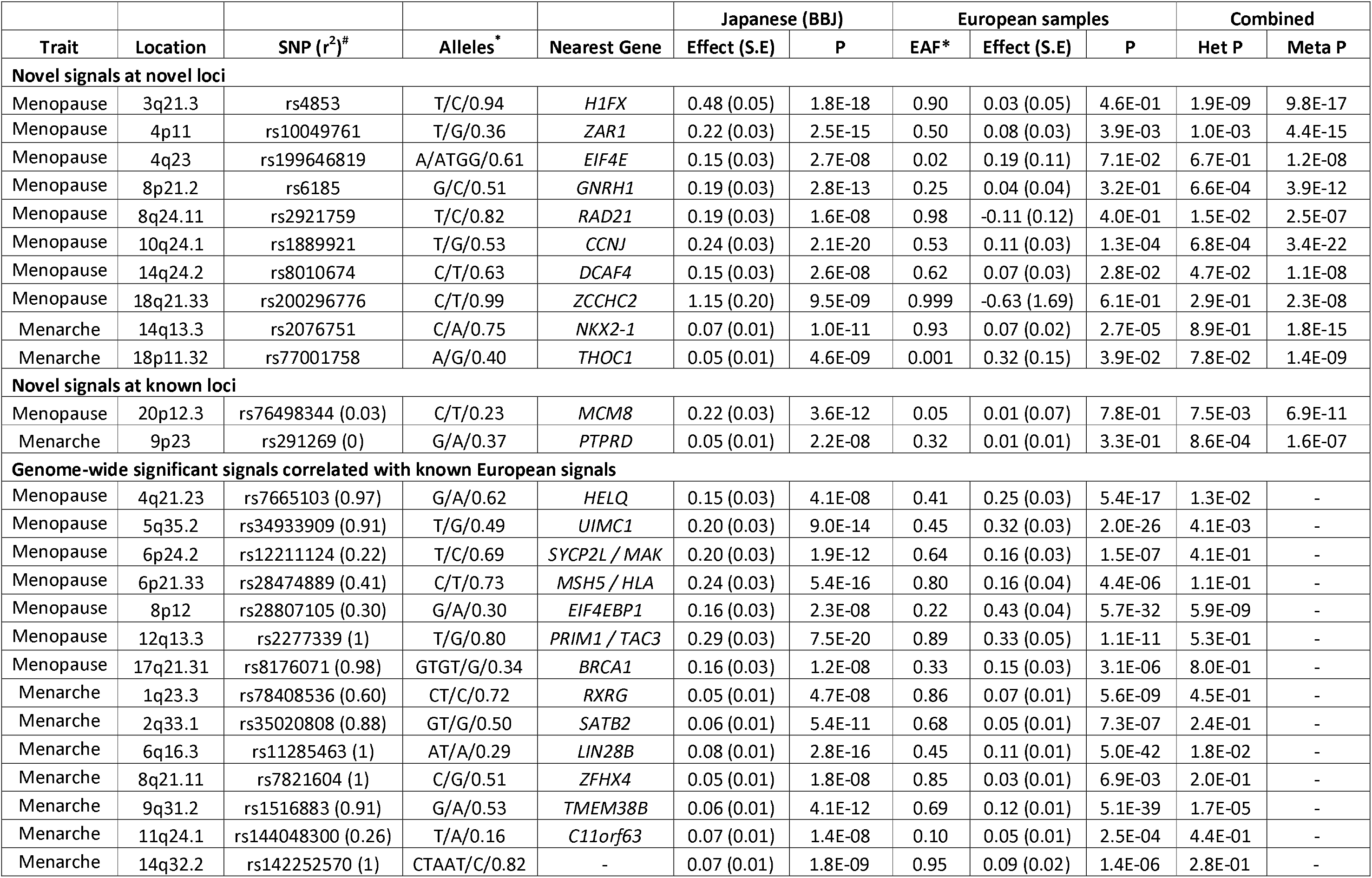
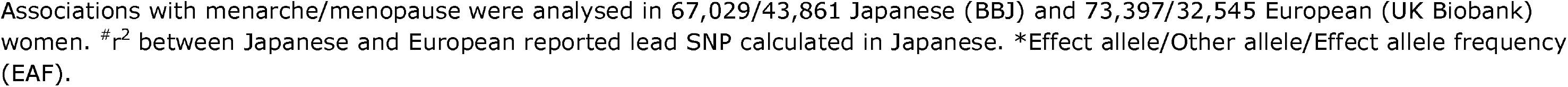
Genome-wide significant signals identified for ages at menarche and menopause in the BioBank Japan Project.

|  |  |  |  |  | Japanese (BBJ) |  | European samples |  |  | Combined |  |
| --- | --- | --- | --- | --- | --- | --- | --- | --- | --- | --- | --- |
| Trait | Location | SNP (r<sup>2</sup>)<sup>#</sup> | Alleles<sup>*</sup> | Nearest Gene | Effect (S.E) | P | EAF* | Effect (S.E) | P | Het P | Meta P |
| Novel signals at novel loci |  |  |  |  |  |  |  |  |  |  |  |
| Menopause | 3q21.3 | rs4853 | T/C/0.94 | H1FX | 0.48 (0.05) | 1.8E-18 | 0.90 | 0.03 (0.05) | 4.6E-01 | 1.9E-09 | 9.8E-17 |
| Menopause | 4p11 | rs10049761 | T/G/0.36 | ZAR1 | 0.22 (0.03) | 2.5E-15 | 0.50 | 0.08 (0.03) | 3.9E-03 | 1.0E-03 | 4.4E-15 |
| Menopause | 4q23 | rs199646819 | A/ATGG/0.61 | EIF4E | 0.15 (0.03) | 2.7E-08 | 0.02 | 0.19 (0.11) | 7.1E-02 | 6.7E-01 | 1.2E-08 |
| Menopause | 8p21.2 | rs6185 | G/C/0.51 | GNRH1 | 0.19 (0.03) | 2.8E-13 | 0.25 | 0.04 (0.04) | 3.2E-01 | 6.6E-04 | 3.9E-12 |
| Menopause | 8q24.11 | rs2921759 | T/C/0.82 | RAD21 | 0.19 (0.03) | 1.6E-08 | 0.98 | -0.11 (0.12) | 4.0E-01 | 1.5E-02 | 2.5E-07 |
| Menopause | 10q24.1 | rs1889921 | T/G/0.53 | CCNJ | 0.24 (0.03) | 2.1E-20 | 0.53 | 0.11 (0.03) | 1.3E-04 | 6.8E-04 | 3.4E-22 |
| Menopause | 14q24.2 | rs8010674 | C/T/0.63 | DCAF4 | 0.15 (0.03) | 2.6E-08 | 0.62 | 0.07 (0.03) | 2.8E-02 | 4.7E-02 | 1.1E-08 |
| Menopause | 18q21.33 | rs200296776 | C/T/0.99 | ZCCHC2 | 1.15 (0.20) | 9.5E-09 | 0.999 | -0.63 (1.69) | 6.1E-01 | 2.9E-01 | 2.3E-08 |
| Menarche | 14q13.3 | rs2076751 | C/A/0.75 | NKX2-1 | 0.07 (0.01) | 1.0E-11 | 0.93 | 0.07 (0.02) | 2.7E-05 | 8.9E-01 | 1.8E-15 |
| Menarche | 18p11.32 | rs77001758 | A/G/0.40 | THOC1 | 0.05 (0.01) | 4.6E-09 | 0.001 | 0.32 (0.15) | 3.9E-02 | 7.8E-02 | 1.4E-09 |
| Novel signals at known loci |  |  |  |  |  |  |  |  |  |  |  |
| Menopause | 20p12.3 | rs76498344 (0.03) | C/T/0.23 | MCM8 | 0.22 (0.03) | 3.6E-12 | 0.05 | 0.01 (0.07) | 7.8E-01 | 7.5E-03 | 6.9E-11 |
| Menarche | 9p23 | rs291269 (0) | G/A/0.37 | PTPRD | 0.05 (0.01) | 2.2E-08 | 0.32 | 0.01 (0.01) | 3.3E-01 | 8.6E-04 | 1.6E-07 |
| Genome-wide significant signals correlated with known European signals |  |  |  |  |  |  |  |  |  |  |  |
| Menopause | 4q21.23 | rs7665103 (0.97) | G/A/0.62 | HELQ | 0.15 (0.03) | 4.1E-08 | 0.41 | 0.25 (0.03) | 5.4E-17 | 1.3E-02 | - |
| Menopause | 5q35.2 | rs34933909 (0.91) | T/G/0.49 | UIMC1 | 0.20 (0.03) | 9.0E-14 | 0.45 | 0.32 (0.03) | 2.0E-26 | 4.1E-03 | - |
| Menopause | 6p24.2 | rs12211124 (0.22) | T/C/0.69 | SYCP2L / MAK | 0.20 (0.03) | 1.9E-12 | 0.64 | 0.16 (0.03) | 1.5E-07 | 4.1E-01 | - |
| Menopause | 6p21.33 | rs28474889 (0.41) | C/T/0.73 | MSH5 / HLA | 0.24 (0.03) | 5.4E-16 | 0.80 | 0.16 (0.04) | 4.4E-06 | 1.1E-01 | - |
| Menopause | 8p12 | rs28807105 (0.30) | G/A/0.30 | EIF4EBP1 | 0.16 (0.03) | 2.3E-08 | 0.22 | 0.43 (0.04) | 5.7E-32 | 5.9E-09 | - |
| Menopause | 12q13.3 | rs2277339 (1) | T/G/0.80 | PRIM1 / TAC3 | 0.29 (0.03) | 7.5E-20 | 0.89 | 0.33 (0.05) | 1.1E-11 | 5.3E-01 | - |
| Menopause | 17q21.31 | rs8176071 (0.98) | GTGT/G/0.34 | BRCA1 | 0.16 (0.03) | 1.2E-08 | 0.33 | 0.15 (0.03) | 3.1E-06 | 8.0E-01 | - |
| Menarche | 1q23.3 | rs78408536 (0.60) | CT/C/0.72 | RXRG | 0.05 (0.01) | 4.7E-08 | 0.86 | 0.07 (0.01) | 5.6E-09 | 4.5E-01 | - |
| Menarche | 2q33.1 | rs35020808 (0.88) | GT/G/0.50 | SATB2 | 0.06 (0.01) | 5.4E-11 | 0.68 | 0.05 (0.01) | 7.3E-07 | 2.4E-01 | - |
| Menarche | 6q16.3 | rs11285463 (1) | AT/A/0.29 | LIN28B | 0.08 (0.01) | 2.8E-16 | 0.45 | 0.11 (0.01) | 5.0E-42 | 1.8E-02 | - |
| Menarche | 8q21.11 | rs7821604 (1) | C/G/0.51 | ZFHX4 | 0.05 (0.01) | 1.8E-08 | 0.85 | 0.03 (0.01) | 6.9E-03 | 2.0E-01 | - |
| Menarche | 9q31.2 | rs1516883 (0.91) | G/A/0.53 | TMEM38B | 0.06 (0.01) | 4.1E-12 | 0.69 | 0.12 (0.01) | 5.1E-39 | 1.7E-05 | - |
| Menarche | 11q24.1 | rs144048300 (0.26) | T/A/0.16 | C11orf63 | 0.07 (0.01) | 1.4E-08 | 0.10 | 0.05 (0.01) | 2.5E-04 | 4.4E-01 | - |
| Menarche | 14q32.2 | rs142252570 (1) | CTAAT/C/0.82 | - | 0.07 (0.01) | 1.8E-09 | 0.95 | 0.09 (0.02) | 1.4E-06 | 2.8E-01 | - |

Associations with menarche/menopause were analysed in 67,029/43,861 Japanese (BBJ) and 73,397/32,545 European (UK Biobank) women. $r^2$ between Japanese and European reported lead SNP calculated in Japanese. \*Effect allele/Other allele/Effect allele frequency (EAF).

Of the 12 novel signals for the two traits, 5 showed larger effect sizes in Japanese than in Europeans (P_heterogeneity_<0.004; i.e. =0.05/12; **Table 1**). A further 3 signals were likely not identified in previous GWAS due to markedly lower allele frequencies in Europeans: *EIF4E*(rs199646819, minor allele frequency (MAF) in Japanese vs. Europeans: 39% vs. 2%), *NKX2-1*(rs2076751: 25% vs. 7%) and *THOC1* (rs77001758: 40% vs. 0.1%). In a meta-analysis allowing for trans-ethnic heterogeneity, 10 of the 12 signals remained genome-wide significant when combined with European data (**Table 1**).

Four novel signals were highly correlated (r^2^>0.7) with missense variants, implicating for the first time the genes *GNRH1, HMCES, ZCCHC2* and *ZNF518A* in the regulation of menopause timing. Notably, rs6185 (Trp16Ser) in *GNRH1* is exactly the same lead SIFT-predicted deleterious missense variant recently reported for age at menarche in Europeans^5^. In our Japanese sample, the rs6185 G-allele was associated with later menopause (beta=0.19 years/allele, P=3×10¯^13^) and later menarche (P=3.5×10¯^5^), but it reportedly has no effect on menopause timing in Europeans (beta=0.03 years/allele, P=0.16, N=67,602). *HMCES* at 3q21.3 encodes an embryonic stem cell-specific binding protein for 5-hydroxymethylcytosine, a recently described epigenetic modification that is dynamically regulated during oocyte ageing^12^, while *ZNF518A* at 10q24.1 encodes an interaction partner of the epigenetic silencing machineries G9a/GLP and Polycomb Repressive Complex 2^13^.

### Genetic associations with early or late menarche timing

As was recently shown in Europeans^5^, we tested in our Japanese GWAS sample whether variants associated with continuous age at menarche have disproportionately larger effects on early versus late puberty timing in females. The approximate earliest (9-12 years inclusive: N = 15,709) and latest (16-20 years: N = 10,875) strata of age at menarche in Japanese were each compared to the same reference group (14 years: N = 14,557). Consistent with findings in Europeans^5^, in Japanese more variants had larger effects on early than on late menarche timing (**Table S5**, 55.7%, 191/343, binomial P=0.02).

We then tested variants genome-wide for early or late menarche timing in Japanese. We identified just one signal at P<5×10¯^8^, rsl0119582 near *PTPRD* associated with early menarche timing (C-allele: OR 1.17 [1.12-1.23], P=8.9×10¯^13^), which was partially correlated with the novel signal for continuous age at menarche in Japanese (r^2^=0.31 with rs291269) and the known European menarche signal at this region (r^2^=0.11 with rs10959016) (**Figure 2A**). Reanalysis of the early menarche model for rs10119582 conditioned on those two other SNPs showed no appreciable change to the magnitude of effect (**Table S6**, beta was attenuated by 13%, conditional P=3.3×10¯^7^). An examination of rs10119582 by each completed whole year of menarche showed that its effects were confined to those ages earlier than the median (age 14), without any apparent effect on menarche timing when older than the median age (**Figure 2B**).

**Figure 2.**
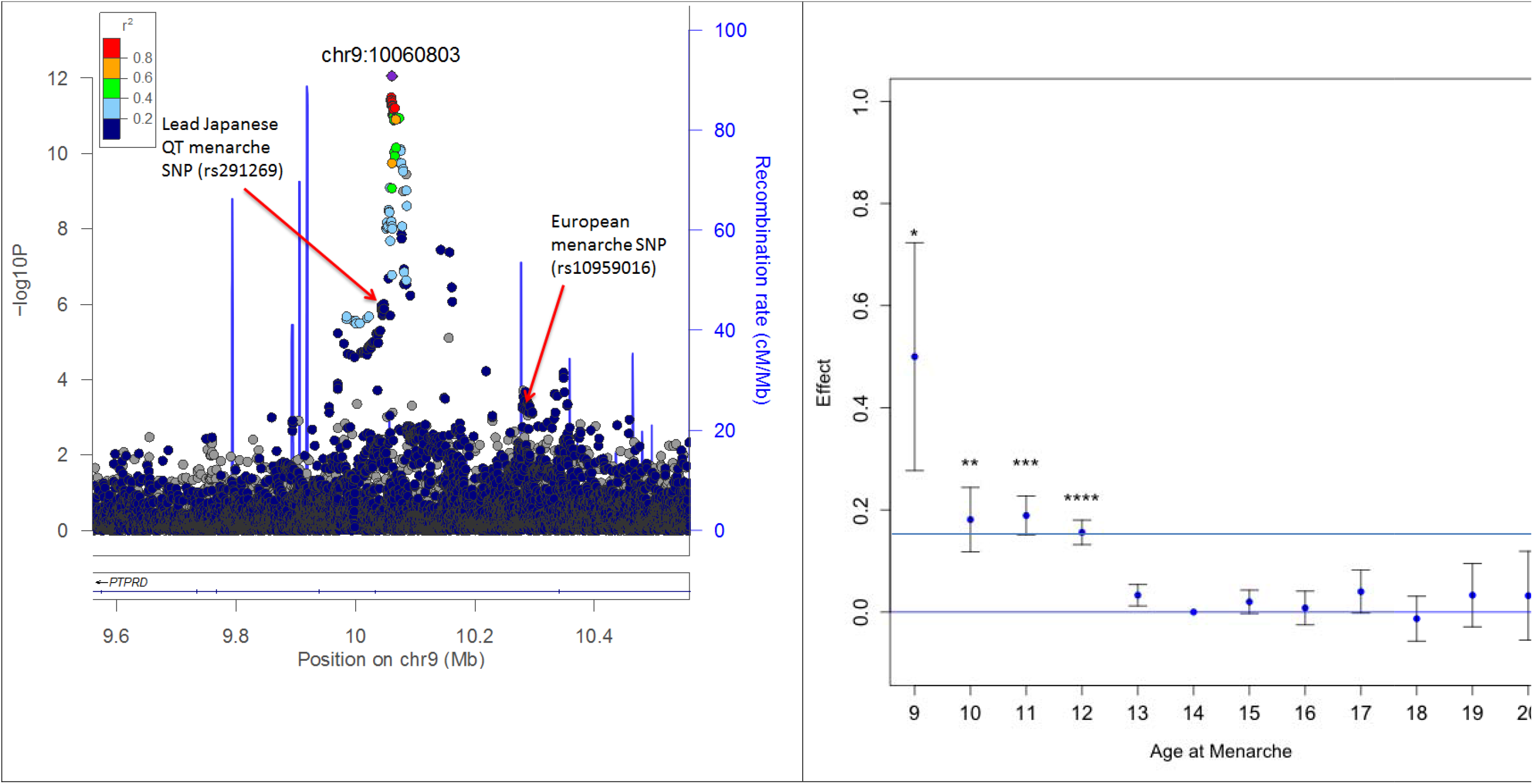
Disproportionate effects on early versus late puberty for rsl0119582 at the *PTPRD* locus. **A:** Regional association plot for the early puberty model in Japanese. **B:**Effect of rsl0119582 at the *PTPRD* locus on risk of specific age; menarche. Each point represents the relative risk (natural log of the odds ratio) per +1 ‘T’ allele at rsl0119582 of being in each menan age category, compared to the reference group (menarche at age 14 years).). *P=0.02, **P=0.004, ***P=6.4×10¯^7^, ****P=1.3×10¯^10^

### Receptor-like protein tyrosine phosphatase genes

Receptor-like tyrosine phosphatases (PTPRs) are a family of 20 cell-surface proteins with intracellular phosphotyrosine phosphatase activity^14^. In addition to the 2 Japanese-specific signals at *PTPRD*, for continuous and early age at menarche, in Europeans 6 further independent signals have been described for menarche (in/near: *PTPRD, PTPRF, PTPRJ, PTPRK, PTPRS*, and *PTPRZ1*) with 2 others just short of genome-wide significance (P<6×10¯^8^, in/near: *PTPRG* and *PTPRN2*). In combination, this gene family is enriched for variant associations with age at menarche (MAGENTA pathway enrichment P_Euro_=7×10¯^3^).

To explore the role of the PTPR gene family in the physiological regulation of puberty timing, we examined changes in gene expression, assessed by RNA-sequencing, in rat medial basal hypothalamus at 5 time points from infancy (postnatal days, PND7 and 14), through juvenile development (EJ, early juvenile PND21; LJ, late juvenile PND28), to the peripubertal period (LP, day of the preovulatory surge of gonadotropins). In false discovery rate-corrected analyses, 13 of the 20 PTPR genes examined were differentially expressed over time: 6 genes were up - regulated (*PTPRB, PTPRC, PTPRJ, PTPRM, PTPRN, PTPRN2*) (all FDR<0.014), and 7 genes were down-regulated (*PTPRD, PTPRG, PTPRK, PTPRO, PTPRS, PTPRT, PTPRZ1*) (Figure 3A % B; Table S7). Additional examination of medial basal hypothalamus expression in a primate model, assessed by quantitative PCR of selected PTPR genes, showed similar time trends in expression as seen in rat hypothalamus: expression of *PTPRN* was up-regulated and *PTPRZ* was down regulated from infancy through puberty (**Figure 3C**).

**Figure 3.**
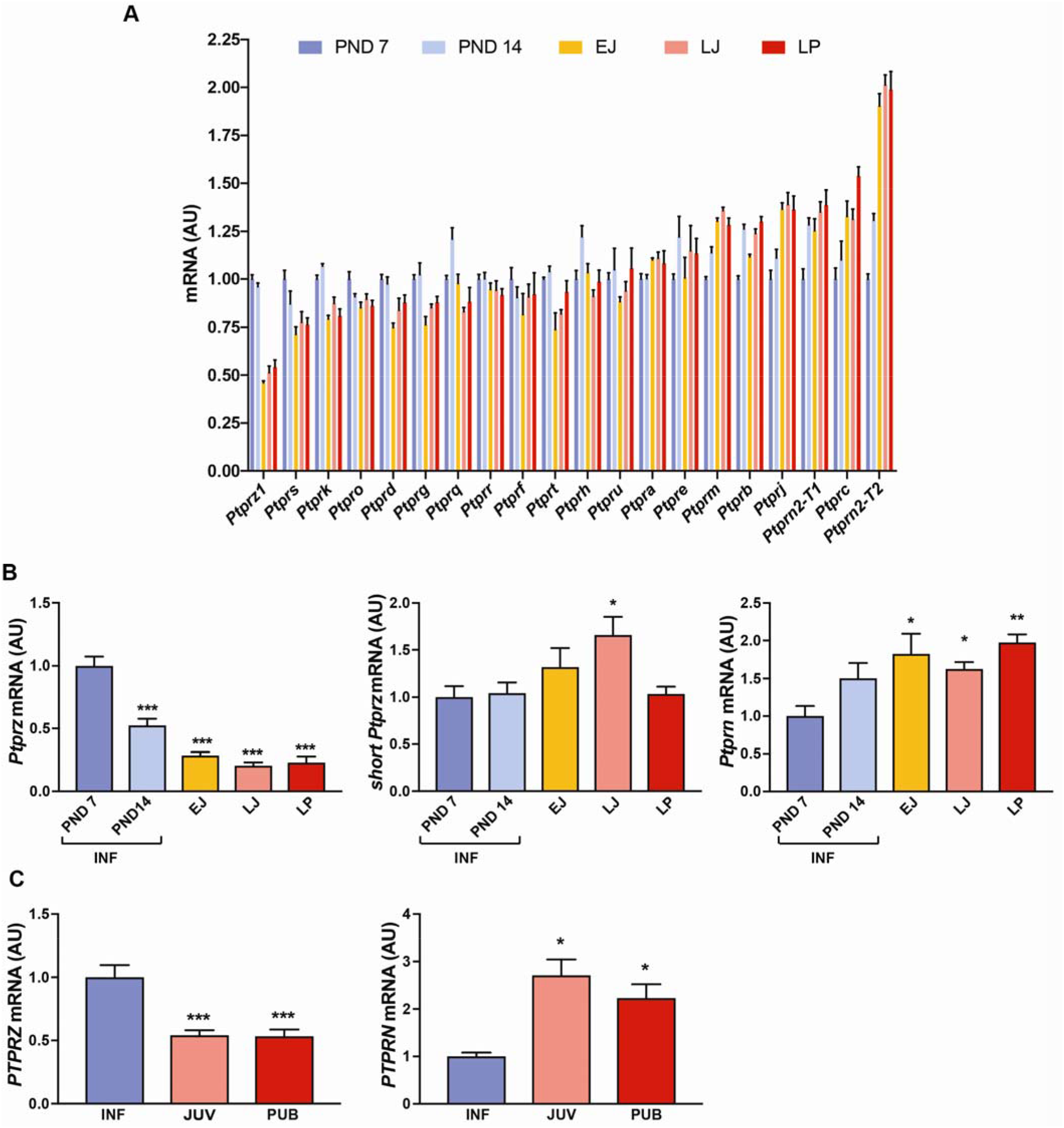
Changes in hypothalamic *PTPR* gene expression during prepubertal development of female rats and rhesus monkeys. (**A**) *PTPR* mRNA levels in the medial hypothalamus (MBH) of pre - and peripubertal female rats assessed by massively parallel sequencing (n=4 biological replicates per developmental stage). (**B**) mRNA abundance of two selected *Ptpr* genes (*Ptprz and Ptprn*) in the MBH of pre - and peripubertal female rats assessed by qPCR. Both the long and short forms of *Ptprz* are shown. Each bar represents the mean ± SEM. *p<0.05; **p<0.01 and ***p<0.001 vs. PND 7, one-way ANOVA followed by Student-Newman-Keuls (SNK) multiple comparison test, n = 6-8 rats per group. INF=infantile period (PND7 and PND14); EJ=early juvenile (PND21); LJ=late juvenile (PND28); LP = late puberty (day of the first preovulatory surge of gonadotropins). (**C**) *PTPRZ* and *PTPRN* mRNA levels in the MBH of pre - and peripubertal female rhesus monkeys. * = p<0.05 and *** = p<0.001 vs. infantile (INF) group, one-way ANOVA followed by SNK multiple comparison test, n= 4-8 monkeys per group. INF= Infantile (1-6 months of age); JUV = Juvenile (6-19 months of age); PUB = Pubertal (24-36 months of age; defined by the presence of elevated plasma LH levels)

## Discussion

Our study represents the largest non-European ancestry genomic analysis for reproductive ageing to date, identifying the first genome-wide significant loci for ages at menarche and menopause outside of Europeans. While the overall heritability estimates were lower in Japanese than in European ancestry populations, we present to our knowledge the first evidence for a secular trend in the heritability for any trait. Secular trends towards earlier age at menarche are widely reported^15^ and are accompanied by secular declines in its variance^16^ - our findings may suggest that such changes may be explained by declining population variability in exposure to environmental factors that delay puberty, such as childhood undernutrition^17^. Despite these population differences in heritability, our findings support a largely shared genetic architecture of reproductive ageing, notably with the replication at genome-wide significance in Japanese of 14 known European signals for menarche or menopause (**Table 1**). However, both effect allele frequencies and effect estimates varied considerably between populations, likely due to a combination of differential genetic drift, selection, recombination and possibly also environmental setting, resulting in substantial heterogeneity in genetic associations between the population groups and reinforcing the need to appropriately model such trans-ethnic differences in meta-analyses.

Such differences in genetic architecture underpin the value of studying genetic associations in diverse population groups to identify novel signals. Hence, even in a Japanese dataset considerably smaller than the largest reported European meta-analysis^5^, we identified ten novel loci for ages at menarche or menopause, and these findings implicated novel genes and pathways as involved in human reproductive ageing. In addition to *HMCES* and *ZNF518A* described above, at the novel menarche locus at 14q13.3, the nearest gene *NKX2-1*, encodes a homeodomain gene that is required for basal forebrain morphogenesis and also remains active in the adult nonhuman primate hypothalamus, where its ablation in mice results in delayed puberty, reduced reproductive capacity, and a short reproductive span^18^, but, until now, has not been implicated in human reproductive function. At the novel menopause locus at 4q23, the nearest gene *EIF4E* encodes a key translation initiation factor; **EIF4E** is the target of the inhibitory binding protein encoded by *EIF4EBP1*, which is near to a known European ancestry menopause locus^7^, while a rare deleterious stop mutation in *EIF4ENIF1*, which encodes a nucleocytoplasmic shuttle protein for **EIF4E**, segregates with primary ovarian insufficiency (menopause at ~30 years old) in a large kindred^19^. Other novel menopause loci implicate for the first time: the evolutionarily conserved maternal-effect gene *ZAR1*, which encodes an oocyte - specific protein that is critical for oocyte-to-embryo transition^20^; *H1FX*, which encodes a member of the histone H1 family; and *RAD21*, a gene involved in chromatid cohesion during mitosis and the repair of **DNA** double-strand breaks, and mutated in two children with Cornelia de Lange syndrome-4, a complex disorder with cellular characteristics of decreased chromatid separation, increased aneuploidy, and defective **DNA** repair^21^.

Our identification of a deleterious variant rs6185 (Trp16Ser) in *GNRH1*, a known signal for age at menarche, as a novel locus for menopause timing suggests an unexpected primary role of hypothalamo-pituitary GnRH signalling in the onset of menopause. Typically, menopause is characterised by ovarian failure and accompanied by a secondary (presumed passive) rise in GnRH-driven gonadotropin secretion. Our finding that the rs6185 G-allele, which delays menarche, also delays menopause is consistent with similar reported effects of alleles near *FSHB*^22,23^, and together suggest that lower levels of gonadotropin secretion may extend reproductive lifespan. Alternatively, GnRH receptor mRNA has been recognized in human ovary, where it may mediate reported autocrine/paracrine actions of GnRH to induce apoptosis of ovarian granulosa cells^24^. Interestingly, despite consistent associations between rs6185 and menarche timing in both Japanese and Europeans, which argues against population differences in LD with an unseen causal variant, we saw a 5-fold greater effect of rs6185 on menopause timing in Japanese than Europeans, a difference that suggests some yet identified strong environmental modification.

Finally, we provide multiple sources of evidence in support of a novel role for receptor-like protein tyrosine phosphatases (PTPRs) in the regulation of puberty timing. The PTPRs are involved in important developmental processes, including the formation of the nervous system by controlling axon growth and guidance^14^. Inactivation of *Ptprs* in the mouse is reported to result in hyposmia and structural defects in the hypothalamus and pituitary^25^. Directly relevant to the regulation of pubertal timing is the observation that in prepubertal female mice a short isoform of PTPRZ1 (also known as RPTPβ) expressed in astrocytes interacts with the glycosylphosphatidyl inositol (GPI)-anchored protein contactin expressed in GnRH neurons^26^. Because contactin is particularly abundant in GnRH nerve terminals, it has been postulated that GnRH neuron-astrocyte communication is in part mediated by RPTPβ-contactin interactions during female reproductive development^26^. In the present study, the implicated PTPR genes are spread across most of the 8 PTPR sub-types (summarised in **Table 2**), indicating that future systematic analyses of the PTPR gene family and potential interactions between these genes would be informative in further understanding the regulation of puberty and related clinical disorders.

**Table 2:** Summary of Receptor-like protein tyrosine phosphatase (PTPR) genes implicated in the regulation of puberty through Japanese and European genome-wide association studies (GWAS) for age at menarche, and pre¯ and peri-pubertal changes in hypothalamic gene expression.

| PTPR sub-type | Genes<sup>1</sup> | GWAS | Expression -Up | Expression -Down |
| --- | --- | --- | --- | --- |
| I | C |  | C |  |
| IIA | D,F,S | D,F,S |  | D,S |
| IIB | K,M,U,T | K | M | K,T |
| III | B,H,J,O,Q | J | B,J | O |
| IV | A,E |  |  |  |
| V | G,Z1 | G,Z1 |  | G,Z1 |
| VI | R |  |  |  |
| VII | N,N2 | N2 | N,N2 |  |

<sup>1</sup>Gene names are abbreviated from *PTPR\**, where \* is a letter (or letter/number).

## Acknowledgements

We would like to acknowledge all the staff in the BBJ project as well as the doctors and comedical staff of the contributing hospitals for their outstanding work on collecting samples and clinical information. We also would like to thank all the patients participating in this project. Work using the UK Biobank Resource was conducted under application 5122.

## Funding

This research was supported by the Tailor-Made Medical Treatment Program (the BioBank Japan Project) of the Ministry of Education, Culture, Sports, Science, and Technology (MEXT) and the Japan Agency for Medical Research and Development (AMED). This work was also supported by the Medical Research Council [Unit Programme number MC_UU_12015/2] and by grants from the US National Science Foundation (NSF: IOS1121691) to S.R.O, and the National Institute of Health (NIH 1R01HD084542) to S.R.O and A.L, and 8P510D011092 for the operation of the Oregon National Primate Research Center. C.A.T was supported by NIH NRSA grant F32-HD-86904. C.A.T and H.W. were supported by NIH Training grants T32-HD007133 and T32-DK 7680. Short read sequencing assays were performed by the OHSU Massively Parallel Sequencing Shared Resource.

## Conflict of interests

The authors declare no conflicts of interests.

## Methods

### BBJ participants and phenotyping

All participants were recruited from Biobank Japan (BBJ), which is a patient-oriented biobank established in Japan^10^. Approximately 200,000 patients diagnosed with any of the 47 targeted common diseases were enrolled in BBJ between 2003 and 2008, and DNA, serum and clinical information were collected from each patient via 66 hospitals across Japan. The current analysis was based on 69,616 female participants who provided information on either age at menarche or menopause and had genotype data available. Ages at menarche or menopause were recalled to the nearest whole completed year at baseline and at multiple follow-up visits. Participants were excluded based on the following conditions: (i) missing age at menarche or menopause; (ii) missing age at recruitment; (iii) maximum difference in the recalled ages at menarche or menopause collected on multiple visits > 5 years; (iv) age at recruitment was younger than reported age at menarche or menopause; (v) missing birth year (for analyses of age at menarche); (vi) age at menarche < 9 or > 20 years; (vii) age at menopause <40 or >60 years and (vii) patients with medical history of hysterectomy, ovariectomy, radiation, chemotherapy and hormone replacement treatment (for analyses of age at menopause). Where age at menarche or menopause was reported at multiple visits, mean values for each were calculated. In total, 67,029 participants with age at menarche and 43,861 with age at menopause were included in the quantitative trait analyses. Age at menarche was also stratified into ‘early’ (ages 9-12 years inclusive, N = 15,709) and ‘late’ (ages 16-20 years inclusive, N = 10,875), and each of these two groups was compared to the same median reference group (age 14, N = 14,557).

### Genotype quality control, imputation and discovery GWAS analysis

BBJ participants had DNA genotyped by either a combination of Illumina Human OmniExpress BeadChip and Infinium HumanExome BeadChip or Infinium OmniExpressExome BeadChip alone. Variants overlapping across these chips were extracted. Variants were then excluded according to the following criteria: (i) monomorphic in any chip; (ii) call rate < 99%; (iii) minor allele count in heterozygotes < 5; (iv) Hardy-Weinberg Equilibrium p-value < 1×10¯^6^ in any chip. For sample quality control, we excluded samples with (i) call rate < 98%; (ii) discordant phenotypic and genotypic sex; (iii) excess heterozygosity; (iv) cryptic relatedness assessed by *pi_hat* measurement (> 0.2) for identity by descent; or if (v) not from mainland Japan identified by principal component analysis using all samples from the 1000 Genomes Project^11^. After quality control, 532,488 autosomal variants were phased using *Eagle2*^27^ and subsequently imputed up to the reference panel from the 1000 Genomes Project Phase 3 using *minimac3*^28^.

Variants with good imputation quality (*minimac* rsq > 0.3)^29^ were tested for associations with two quantitative traits, age at menarche and age at menopause, and two dichotomous traits, early menarche and late menarche, assuming additive allelic effects. Associations with ages at menarche and menopause were tested in linear regression models using *mach2qtl*^30^.

Associations with early and late menarche (both versus the median group), or each age year of age at menarche (age 9, 10, 11, 12, 13, 15, 16, 17, 18, 19, 20) versus the same median (age 14) group), were tested in logistic regression models using *mach2dat*^30^. In each model, ten principal components were included to adjust for cryptic population structure, in addition to birth year as a covariate.

Variance explained by genetic variants in the current study were estimated using the *REML* method implemented in *BOLT-LMM*^31^. We tested different SNP sets: (i) Directly genotyped variants within 250 kb up - or down-stream of the previously reported European lead variants^5^’^7^, and (ii) all directly genotyped variants which passed quality control. Pathway enrichment for PTP receptor genes was assessed using *MAGENTA*^32^ on the most recently reported European GWAS meta-analysis.

### Effect estimate comparisons with samples of European ancestry

Known menarche and menopause European loci were defined as those discovered in the two largest reported GWAS meta-analyses to date^5,7^. As effect estimates reported in discovery metaanalyses are potentially inflated due to the ‘winners curse’ phenomenon, we derived more robust European effect estimates in independent samples from the UK Biobank study^33^. For menarche, this required us to restrict the number of known European loci to the largest discovery meta-analysis prior to inclusion of UK Biobank^5^. A total of 73,397 women with genotype and age at menarche were available from UK Biobank, and analysis of this sample has been described previously^5^. Age of natural menopause was available for 32,545 UK Biobank women, using the same inclusion/exclusion criteria applied to BBJ women. This analysis was performed using a linear mixed model implemented in *BOLT*, as previously described^34^. Transethnic meta-analysis was performed using Han and Eskin’s Random Effects model, implemented in *Metasoft*^35^.

### Animal samples for hypothalamic gene expression

Sprague Dawley female rats were studied at different phases of postnatal development: infantile postnatal day (PND) 7 and 14, early juvenile (EJ) PND21, late juvenile (LJ) PND28, and late puberty (LP, the day of the first preovulatory surge of gonadotropins, PND32-38. The use of rats was approved by the ONPRC Animal Care and Use Committee in accordance with the NIH guidelines for the use of animals in research. The animals were obtained from Charles River Laboratories international, Inc. (Hollister, CA), and were housed in a room with controlled photoperiod (12/12 h light/dark cycle) and temperature (23-25°C). They were allowed *ad libitum* access to pelleted rat chow and water. The medial basal hypothalamus (MBH) of female rats was collected at various postnatal ages by performing a rostral cut along the posterior border of the optic chiasm, a caudal cut immediately in front of the mammillary bodies, and two lateral cuts half-way between the medial eminence and the hypothalamic sulci. The thickness of the resulting tissue fragment was about 2 mm. The fragment includes the entire arcuate nucleus (ARC). Upon dissection, the tissues were immediately frozen on dry ice and stored at -85°C until RNA extraction.

Female rhesus monkey (*Macaca mulatta*) hypothalamic tissue samples were obtained through the Oregon National Primate Research Center (ONPRC) Tissue Distribution Program for the studies of infantile-pubertal (INF-PUB) transitions. The animals were classified into different stages of development based on their age and pubertal stages, following the recommendation reported by Watanabe and Terasawa^36^. The brain was removed from the cranium and the MBH was dissected as previously described^37^, that is, by making a rostral cut along the posterior border of the optic chiasm, a caudal cut immediately in front of the mammillary bodies, and two lateral cuts half-way between the medial eminence and the hypothalamic sulci. The tissue fragments were rapidly frozen by immersion in liquid nitrogen and stored at -80°C.

### RNA extraction and reverse transcription PCR

Total RNA was extracted from the MBH tissues of female rats and rhesus monkeys at different developmental stages using the RNeasy mini kit (Qiagen, Valencia, CA) following the manufacturer’s protocol. DNA contamination was removed from the RNA samples by on-column digestion with DNAse using the Qiagen RNase-free DNase kit (Qiagen, Valencia, CA) according to the manufacturer’s instructions. RNA concentrations were determined by spectrophotometric trace (Nanodrop, ThermoScientific, Wilmington, DE). Five-hundred ng of total RNA were transcribed into cDNA in a volume of 20 µl using 4 U of Omniscript reverse transcriptase (Qiagen, Valencia, CA).

Relative mRNA abundance was determined using the SYBR GreenER™ qPCR SuperMix system (Invitrogen, Carlsbad, CA). Suitable amplification primers were designed using the PrimerSelect tool of DNASTAR 14 software (Madison, WI) or the NCBI online Primer-Blast program (**Table S8**). PCR reactions were performed in a volume of 10 µl containing 1 µl of diluted cDNA, 5 µl of SYBR GreenER™ qPCR SuperMix and 4 µl of primers mix (1 µM of each gene specific primer). The PCR conditions used were as follows: 5 min at 95°C, 40 cycles of 15 sec at 95°C and 60 sec at 60°C. To confirm the formation of a single SYBR Green-labeled PCR amplicon, each PCR reaction was followed by a three-step melting curve analysis consisting of 15 sec at 95°C, 1 min at 60°C, ramping up to 95°C at 0.5°C/sec, detecting every 0.5 sec and finishing for 15 sec at 95°C, as recommended by the manufacturer. All qPCR reactions were performed using a QuantStudio 12K Real-Time PCR system (Thermo Fisher, Waltham, MA); threshold cycles (CTs) were detected by QuantStudio 12K Flex software (Thermo Fisher, Waltham, MA). Relative standard curves were constructed from serial dilutions (1/2 to 1/512) of a pool of cDNAs generated by mixing equal amounts of cDNA from each sample. The CTs from each sample were referred to the relative standard curve to estimate the mRNA content/sample; the values obtained were normalized for procedural losses using glyceraldehyde-3-phosphate dehydrogenase (*GAPDH*) mRNA as the normalizing unit.

### Next generation RNA sequencing (RNA-seq) and analysis

Total RNA obtained from the MBH of female rats at different stages of prepubertal development was subjected to RNA-seq. The procedure was carried out by the OHSU Massively Parallel Sequencing Shared Resource. RNA-seq libraries were prepared using a TruSeq Stranded protocol with ribosomal reduction (Illumina, San Diego, CA). In brief, 600 ng of total RNA per sample were depleted of ribosomal RNA using RiboZero capture probes (Illumina, San Diego, CA). Later, the purified RNA was fragmented using divalent cations and heat, and was used as template for reverse transcription using random hexamer primers. The resulting cDNAs were then treated enzymatically to generate blunt ends. Thereafter, a single “A” nucleotide was added to the 3’ ends to facilitate adaptor ligation. Standard six-base pair Illumina adaptors were ligated to the cDNAs and the resulting DNA was amplified by 12 rounds of PCR. All procedures were carried out following the protocol provided by Illumina. Unincorporated material was removed using AMPure XP beads (BeckmanCoulter, Brea, CA). Libraries were profiled on a Bioanalyzer instrument (Agilent, Santa Clara, CA) to verify the distribution of DNA sizes and the absence of adapter dimers. Library titers were determined by real time PCR (Kapa Biosystems, Wilmington, MA) using a StepOnePlus Real Time System (ThermoFisher, Waltham, MA). Libraries were mixed to run four samples per lane on the HiSeq 2500 (Illumina). Sequencing was carried out using a single-read 100-cycle protocol. The resulting base call files (.bcl) were converted to standard fastq formatted sequence files using Bcl2Fastq (Illumina). Sequencing quality was assessed using FastQC (Babraham Bioinformatics, Cambridge, UK). The RNA-seq data was deposited in NCBI under the accession number GSE94080.

The differential expression of genes during pubertal development was determined by employing the gene-level edgeR^38^ analysis package. An initial trimming and adapter removal step was carried out using Trimmomatic^39^. The reads that passed the Trimmomatic selection criteria were then aligned to the rn6 build of the rat genome with Bowtie2/Tophat2^40,41^, and assigned to gene-level genomic features with the Rsubread featureCounts package based on the Ensembl 83 annotation set. Differential expression between time points was analyzed using the generalized linear modeling approaches implemented in edgeR. Batch effect terms were included in these models to correct for runs on different dates/flow cells. Differentially expressed genes/transcripts were identified based on significance of pairwise comparison of time points to identify the genes most likely to be differentially expressed for later RT-qPCR confirmation.

All statistical analyses were performed using Prism7 software (Graphpad Software, La jolla, CA). The differences between groups were analyzed by ONE WAY ANOVA followed by the Student-Newman-Keuls multiple comparison test for unequal replications. When comparing percentages, groups were subjected to arc-sine transformation before statistical analysis to convert them from a binomial to a normal distribution. A p value of < 0.05 was considered statistically significant.

